# Elucidation of the glycan structure of the b-type flagellin of *Pseudomonas aeruginosa* PAO1

**DOI:** 10.1101/2024.09.23.614423

**Authors:** Paul J. Hensbergen, Loes van Huijkelom, Jordy van Angeren, Arnoud H. de Ru, Bart Claushuis, Peter A. van Veelen, Wiep Klaas Smits, Jeroen Corver

**Author notes:** Correspondence to: P.J. Hensbergen, Center for Proteomics and Metabolomics, Leiden University Medical Center, PO Box 9600, 2300 RC Leiden, The Netherlands.

## Abstract

Flagella are essential for motility and pathogenicity in many bacteria. The main component of the flagellar filament, flagellin, often undergoes post-translational modifications, with glycosylation being a common occurrence. In *Pseudomonas aeruginosa* PAO1, the b-type flagellin is *O*-glycosylated with a structure that includes a rhamnose, a phospho-group and a previous unknown moiety. This structure resembles the well-characterized glycan (Type A) in *Clostridioides difficile* strain 630, which features an *N*-acetylglucosamine linked to an *N*-methylthreonine via a phosphodiester bond.

This study aimed to characterize the b-type glycan structure in *Pseudomonas aeruginosa* PAO1 using a set of mass spectrometry experiments. For this purpose, we used wildtype *P. aeruginosa* PAO1 and several gene mutants from the b-type glycan biosynthetic cluster. Moreover, we compared the mass spectrometry characteristics of the b-type glycan with those of *in vitro* modified Type A-peptides from *C. difficile* strain 630Δ*erm*.

Our results demonstrate that the thus far unknown moiety of the b-type glycan in *P. aeruginosa* consists of an *N,N*-dimethylthreonine. These data allowed us to refine our model of the flagellin glycan biosynthetic pathway in both *P. aeruginosa* PAO1 and *C. difficile* strain 630.

## Introduction

Bacterial flagella are intricate, whip-like appendages that extend from the cell bodies of many motile bacteria, playing a crucial role in their locomotion and environmental navigation (1). These structures are not only essential for bacterial motility but also contribute significantly to pathogenicity, colonization, and biofilm formation (2, 3). At the core of the flagellar filament is FliC, also known as flagellin, a highly conserved protein across many bacterial species that polymerizes to form the helical structure driving bacterial movement (4).

Beyond its structural role, FliC undergoes various post-translational modifications (5), among which glycosylation is particularly important (6). Glycosylation of FliC can affect the assembly and function of flagella, influencing the stability, flexibility, and overall performance of the flagellar filament. This modification is not uniform across bacterial species; different bacteria employ distinct glycosylation patterns, which can affect their motility in various ways (7, 8). Moreover, FliC glycan structures can vary between different strains of the same species.

Flagellin glycosylation is also observed in *P. aeruginosa,* an aerobic, gram-negative bacterium that primarily causes infections in immunocompromised individuals or patients receiving intensive care (9, 10). Importantly, *P. aeruginosa* strains without flagellin glycosylation showed attenuated virulence (11). In the *P. aeruginosa* PAK strain, the a-type flagellin is *O*-glycosylated with a structure comprising 11 monosaccharide residues (12, 13). In contrast, the b-type flagellin, as observed in the *P. aeruginosa* PAO1 strain, is decorated with a very different structure **Fig**. **1A**) that consists of a single *O*-linked deoxyhexose, likely a L-rhamnose (14), linked to an unknown moiety through a phosphodiester bond (15). Initial experiments indicated that the unknown moiety consists of a tyrosine (16), however subsequent studies by mass spectrometry (15) determined a mass for this moiety (129 Da) which is not compatible with a tyrosine. Hence, the nature of the unknown moiety has been a longstanding unsolved question.

**Figure 1:**
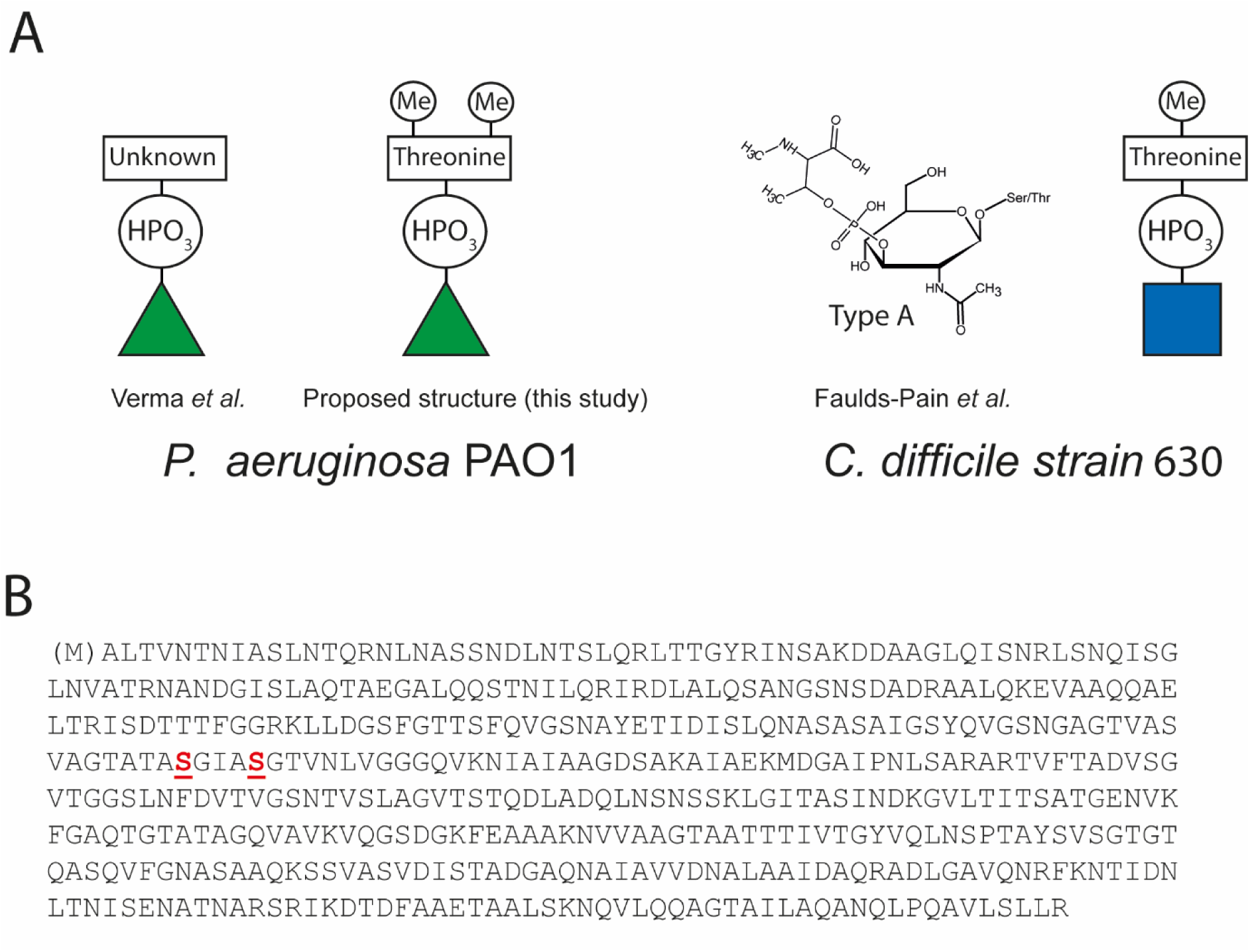
Flagellin glycan structures in *Pseudomonas aeruginosa* PAO1 and *Clostridioides difficile* strain 630. A. Schematic structure of the described ((15)) and proposed structure of the b-type glycan in *P. aeruginosa* and the molecular and schematic structure of the Type A glycan in C. difficile strain 630 ((17)). 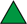: rhamnose (based on (14)). 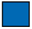: N-acetylglucosamine. HPO_3_: phospho. Me: methyl. **B**. Primary sequence of b-type flagellin of *P. aeruginosa* PAO1 (FliC, UniprotID: P72151). The glycosylated serine residues as determined by Verma et al. (15) are highlighted in red, and underlined. The N-terminal methionine is lacking in the mature protein.

Interestingly, the glycan structure in *P. aeruginosa* has overlapping features to a glycan structure which is found on FliC in several *Clostridioides difficile* strains. This glycan structure (Type A, **Fig. 1A**) consists of an *O*-linked *N*-acetylglucosamine (GlcNAc), that is linked to *N*-methyl-L-threonine through a phosphodiester bond (17, 18). In support of the structural similarity between the two species, gene clusters encoding enzymes with similar activities are found in both genomes (**Fig. S1**) (15, 17). Based on our recent experiments with *C. difficile* strain 630Δ*erm* and comparative bioinformatics of the *P. aeruginosa* and *C. difficile* gene clusters, we have hypothesized that the unknown moiety in the b-type FliC glycan structure in *P. aeruginosa* is an *N,N*-dimethylthreonine (19, 20) **Fig. 1A**). Here, using mass spectrometry-based analyses, we provide strong evidence supporting the presence of this structure on *P. aeruginosa* PAO1 flagellin.

## Results

### Mass spectrometric analysis of b-type glycan-modified flagellin of P. aeruginosa PAO1 is consistent with the presence of a N,N dimethylthreonine as part of the glycan structure

A previous study showed that b-type flagellin from *P. aeruginosa* PAO1 is glycosylated at Ser-191 and Ser-195 in the mature protein (15), **Fig. 1B**)). For these experiments, purified flagella were used. To study the *P. aeruginosa* PAO1 flagellin glycosylation, we sought to apply a phosphoproteomics approach (Immobilized Metal Affinity Chromatography (IMAC)) that we recently used to enrich for Type A-modified tryptic FliC peptides from *C. difficile* (20). Given the similarity in the glycan structures between *C. difficile* strain 630 and *P. aeruginosa* PAO1, most importantly the presence of the phospho-moiety Fig. **1A**), we hypothesized that this approach would also work to enrich for b-type glycan-modified peptides from *P. aeruginosa* PAO1. A Proteinase K digest of the *P. aeruginosa* PAO1 proteome was performed to generate small peptides that would allow us to pinpoint the modified residues. Following IMAC affinity purification, peptides were analysed by LC-MS/MS. The most intense peaks in the LC-MS/MS data represented various b-type glycan-modified peptides originating from flagellin (Fig. **2A**).

**Figure 2:**
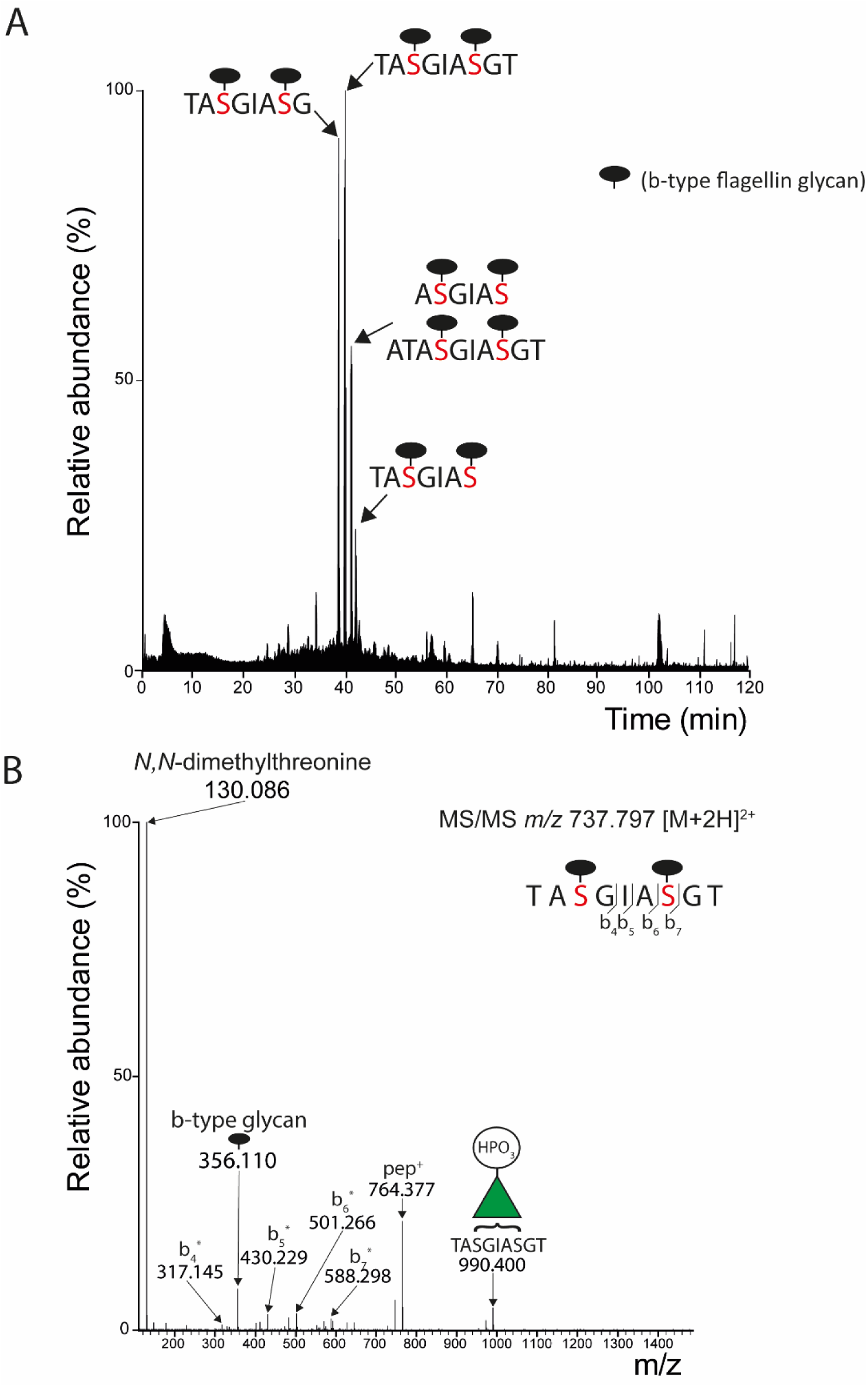
Mass spectrometry analysis of b-type glycan-modified peptides from *P. aeruginosa* PAO1 after IMAC purification. A: Total ion chromatogram of an LC-MS/MS analysis of IMAC-purified Proteinase K peptides from *P. aeruginosa* PAO1. The major b-type glycan-modified peptides corresponding to the observed peaks are indicated. See also Table 1. B: MS/MS spectrum of the Proteinase K flagellin peptide TASGIASGT, modified with two type-b glycans. All observed b-ions have lost the b-type glycan (indicated with a *).

**Table 1:**
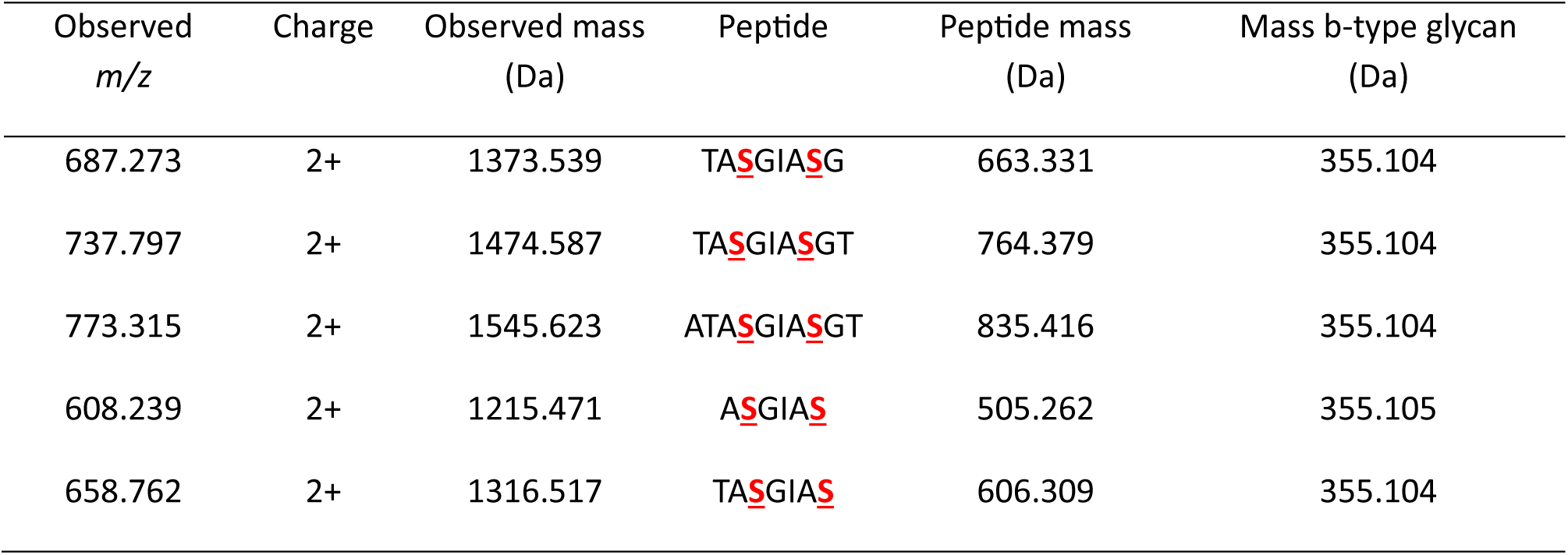
*Probing the b-type glycan based on the MS analysis of Proteinase K generated b-type glycan-modified flagellin peptides from* P. aeruginosa. On each peptide, two b-type glycans were found (serines indicated in red and underlined). See also Figure 2.

The smallest peptide (ASGIAS) allowed us to confirm that Ser-191 and Ser-195 of flagellin are modified with the b-type glycan. Based on the mass of the five major glycopeptides (**Fig. 2A**), a neutral mass of 355.104 Da for the b-type glycan was observed (**Table 1**).

These results are consistent with our hypothesis that the unknown moiety is a *N,N*-dimethylthreonine (theoretical neutral mass of predicted b-type glycan is 355.103 Da (**Fig. 1A**)). MS/MS analysis of the peptide TASGIASGT with two b-type glycans (**Fig. 2B**) showed the loss of the full type-b glycan at *m/z* 356.110 ([M+H^+^]^+^). The MS/MS data also showed a prominent ion at *m/z* 130.086 ([M+H^+^]^+^) (**Fig. 2B**). In line with the proposed b-type glycan structure (**Fig. 1A**), this ion was tentatively assigned as *N,N*-dimethylthreonine. Of note, MS3 experiments showed that the ion at *m/z* 130.086 was derived from the b-type glycan (**Fig. S2**).

### P. aeruginosa PAO1 pa1088-pa1091 mutant strains exhibit truncated b-type glycan structures

In *C. difficile* strain 630, the biosynthesis of the Type A glycan depends on *cd0240* (encoding a glycosyltransferase) and the adjacent *cd0241-cd0244* operon (17). In the *P. aeruginosa* PAO1 biosynthetic gene cluster (**Fig. S1**), only four genes are found (*pa1088*-*pa1091*) because the glycosyltransferase and *cd0244* homolog are organized in a single gene (*pa1091*). To study the role of the individual genes, we recently analyzed the Type A glycan structure and variations thereof in *C. difficile* strain 630 *cd0241-cd0244* mutant strains using a quantitative proteomics experiment. In the *cd0241, cd0242* and *cd0244* mutant strains, truncated Type A structures consisting of only the core GlcNAc were observed. In the *cd0243* mutant, Type A structures solely lacking the methyl group were found, which is in line with the putative methyltransferase activity of CD0243. The methyltransferase homolog in *Pseudomonas aeruginosa* PAO1 is *pa1088* (**Fig. S1**). Hence, based on our proposed b-type glycan structure (**Fig. 1A**), we predicted a truncated b-type glycan lacking two methyl groups in a *pa1088* mutant strain. To test this hypothesis, we performed a quantitative proteomics experiment using *Pseudomonas aeruginosa* PAO1 wildtype (WT) and *pa1088-pa1091* mutant strains. To increase the homogeneity in peptides, we used a combination of trypsin and chymotrypsin instead of Proteinase K for these experiments. All strains were analyzed in triplicate, resulting in a TMT-15 plex experiment.

First, the sample was affinity purified using IMAC. The digestion protocol resulted in the tryptic+chymotryptic peptide QVGSNGAGTVASVAGTATASGIASGTVNLVGGGQVK (aa 172-207) carrying two b-type glycans.

This peptide was observed at *m/z* 1116.317 ([M+4H^+^]^4+^. Again, the MS/MS spectrum **Fig. 3A**) showed the b-type glycan-specific ions at *m/z* 130.086 and 356.110. Zooming in on the TMT-reporter region (**Fig. 3A** (inset) and **Fig. 3B** (upper left panel)) showed that this peptide was observed in the WT samples (TMT-reporter 126, 127N and 127C) but not in the mutant strains. Next, we looked for the same peptide with b-type glycans lacking two methyl groups, which we would expect in the *pa1088* mutant strain, but such a peptide was not observed. Also, peptides with b-type glycans lacking one methyl group were not identified. As expected, given the IMAC affinity purification, mutants with truncated structures lacking the phospho-moiety, e.g. non-glycosylated or only consisting of the rhamnoses, were not observed. To test whether such peptides were present in the *pa1088* mutant and the other strains, we also analysed the sample without the IMAC affinity purification step. For this purpose, the full TMT-labeled digest was fractionated in twelve fractions using high-pH reversed phase chromatography, and each fraction was analyzed by LC-MS/MS.

**Figure 3:**
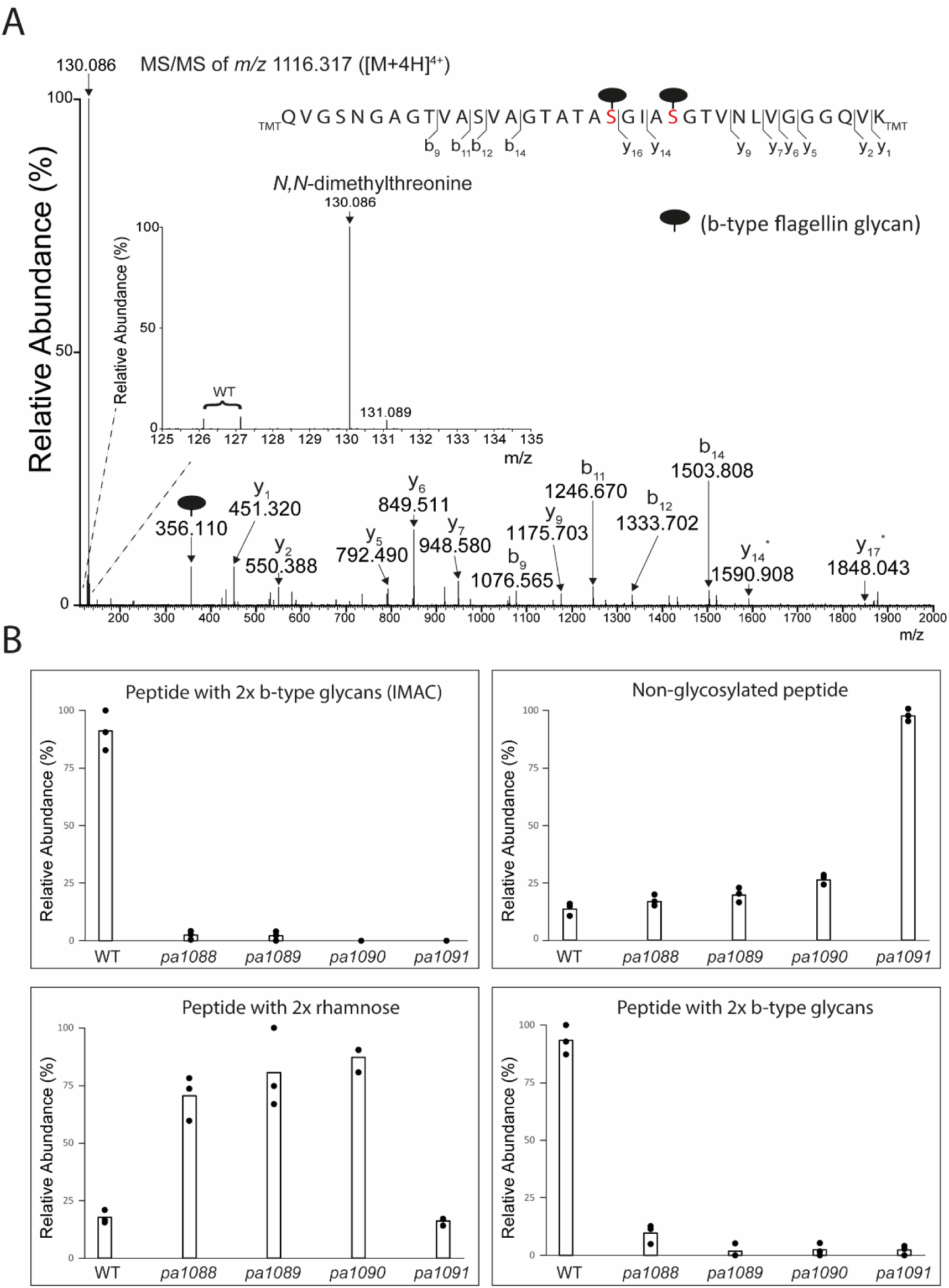
Analysis of variations of the b-type glycan structure in P. aeruginosa PAO1 WT and mutant strains. A: Protein extracts of WT and pa1088-pa1091 *P. aeruginosa* PAO1 (mutant) strains were digested with a combination of trypsin and chymotrypsin. All strains were analyzed in triplicate. Following TMT-labeling, the sample was affinity purified using IMAC and analysed by LC-MS/MS. Shown is the MS/MS spectrum of the peptide QVGSNGAGTVASVAGTATASGIASGTVNLVGGGQVK with two b-type glycan structures. The inset depicts the zoom-in of the TMT-reporter region, which also shows the b-type glycan-specific fragment at m/z 130.086. Fragment ions indicated with an * have lost the b-type glycans. B: Quantification of the different variants of the QVGSNGAGTVASVAGTATASGIASGTVNLVGGGQVK flagellin peptide in WT and individual *P. aeruginosa* PAO 1 mutant strains, i.e., carrying two b-type glycans (top left panel (IMAC purified) and lower right panel (without IMAC), non-glycosylated (top right panel) and carrying two rhamnoses (lower left panel). Quantification was based on the relative intensity of the corresponding TMT-reporter ions. Dots indicate the relative intensity for each biological replicate.

First, we checked our data for the presence of PA1088-PA1091 to explore if we could confirm the mutant phenotype. For PA1088-PA1090, only a few (quantifiable) peptides were observed and the quantitative information based on the relative intensity of the reporter ions was not consistent with the mutant phenotype, even though the data indicated lower levels of PA1088 and PA1090 in the corresponding mutant strains (**Table S3**). We observed a higher protein protein coverage for PA1091 but the quantitative proteomics data could not confirm the mutant phenotype (**Table S3**). However, there appeared to be a difference in peptides covering the N- and C-terminal regions of PA1091. Compared to WT, peptides covering the C-terminal region were found at lower levels, while peptides covering the N-terminal region were not (**Fig. S3**). Interestingly, this correlates with the position where the transposon is integrated into the *pa1091* gene in this strain. This indicates that the N-terminus is still translated, but this does not result in an active protein.

Next, we checked the data for the different variations in the b-type glycan on the tryptic+chymotryptic peptide QVGSNGAGTVASVAGTATASGIASGTVNLVGGGQVK, i.e., the non-glycosylated peptide and the peptide with truncated b-type glycan consisting of only the core rhamnoses. Both were identified with MS/MS spectra that look very different from the b-type glycan-modified peptides because they lacked the prominent b-type glycan-specific ions at *m/z* 130.086 and 356.110 (**Fig. S4**). On the other hand, the peptide-specific fragments largely overlapped. Based on the TMT-reporter intensities in these spectra, the non-glycosylated peptide was predominantly observed in the *pa1091* mutant strain (**Fig. 3B**, upper right panel). In contrast, the peptide with a truncated b-type glycan consisting of only the core rhamnoses was enriched in the *pa1088*, *pa1089*, and *pa1090* mutant strains (**Fig**. **3B**, lower left panel). The fully glycosylated peptide was also identified in the overall proteomics dataset and the quantification data (**Fig. 3B**, lower right panel) agreed with the data from the IMAC purified material (**Fig**. **3B**, upper left panel). The peptide with b-type glycans lacking one or two methyl groups was also not observed in the overall proteomics data.

Collectively, our data show that insertional mutagenesis of genes *pa1088*-*pa1091* in *P. aeruginosa* PAO1 leads to aberrant glycosylation of b-type flagellin. In three mutants, i.e., *pa1088*-*pa1090*, only rhamnoses were observed, while non-glycosylated flagellin was found in the *pa1091* mutant strain.

### Tandem mass spectrometry-based assignment of N,N-dimethylthreonine as part of the b-type glycan in *P. aeruginosa*

Because the *pa1088* mutant data shown above did not demonstrate b-type glycan structures lacking methyl groups, we sought an alternative approach to substantiate our assignment of the *N,N*- dimethylthreonine as part of the b-type glycan. We reasoned that *in vitro* methylation of the Type A structure in *C. difficile* should generate a glycan structure identical to our proposed b-type glycan in *P. aeruginosa*, except for the monosaccharide (**Fig. 1A**). We hypothesized that the mass spectrometric fragmentation characteristics of both structures should be highly similar.

To test this, we methylated tryptic peptides from a WT *C. difficile* 630Δ*erm* strain by reductive amination. With this protocol, all peptide N-termini and lysine side chains were di-methylated. The Type A glycan already contains an *N*-methylated threonine (**Fig. 1A**); therefore, only one methyl group was added resulting in a modified Type A structure with two methyl groups.

During the LC-MS/MS analysis, we focussed on one of the tryptic FliC peptides from *C. difficile* (LLDGTSSTIR) with the methylated Type A structure (**Fig. 4A**). Besides the peptide-specific fragments, a few differences with a b-type glycan-modified peptide from *P. aeruginosa* were apparent in the MS/MS data. First of all, the loss of the fully methylated Type A moiety, which would result in a fragment at *m/z* 413.132 ([M+H^+^]^+^) was hardly observed (**Fig. 4A**). Instead, partial fragmentation of the methylated Type A was apparent. For example, a fragment at *m/z* 228.063 ([M+H^+^]^+^) corresponding to an *N,N*-dimethylthreonine-phosphate was observed. The corresponding fragment at *m/z* 214.048, lacking one methyl group, is well known from the fragmentation spectra of WT Type A-modified peptides (18, 20). Moreover, partial fragmentation of the methylated Type A was observed by fragments at e.g., *m/z* 1275.677, 1293.687 and 1373.654.

**Figure 4:**
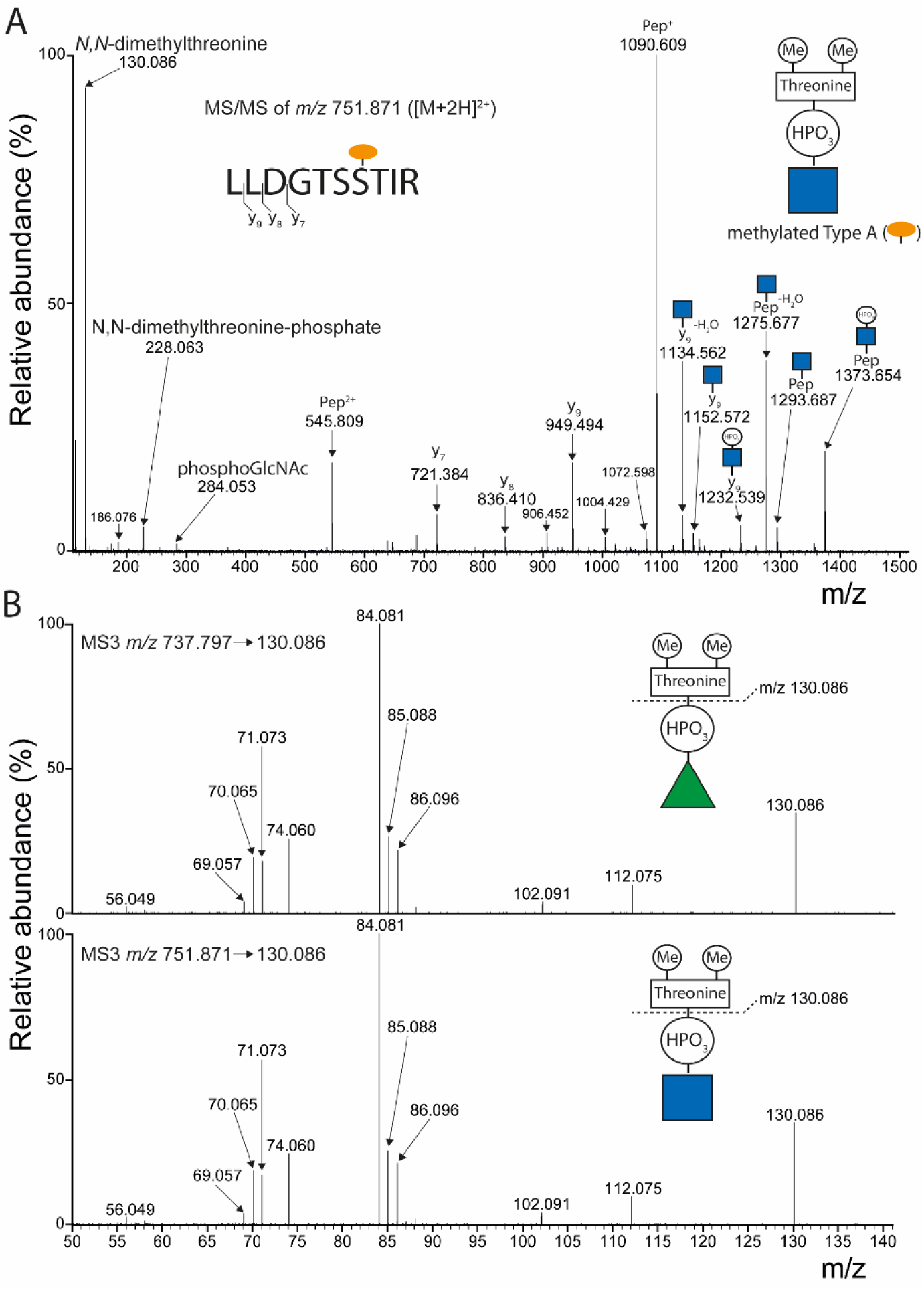
Mass spectrometric identification of *N,N*-dimethylthreonine as part of the b-type glycan in *P. aeruginosa* PAO1. A: A tryptic digest of C. difficile strain 630 Δerm proteins was in vitro methylated using reductive amination and analyzed by LC-MS/MS. Shown is the MS/MS spectrum of the methylated Type A-modified tryptic flagellin peptide LLDGTSSTIR (site assignment based on (17)). The fragment ion at m/z 130.086 corresponds to N,N-dimethylthreonine. B: Comparison of the fragmentation pattern of the ion at m/z 130.086 generated by fragmentation of a *P. aeruginosa* type-b modified peptide and a methylated Type A-modified C. difficile peptide, respectively. Upper panel: Fragmentation (MS3) of the ion at m/z 130.086 generated by MS/MS of the b-type glycan modified peptide TASGIASGT from *P. aeruginosa* flagellin (see Fig. 2B). Lower panel: Fragmentation (MS3) of the ion at m/z 130.086 generated by MS/MS of the methylated Type A-modified peptide LLDGTSSTIR from C. difficile flagellin (see panel A).

Most importantly, the MS/MS spectrum showed the prominent ion at *m/z* 130.086 ([M+H^+^]^+^). In this case, it could be confidently assigned as corresponding to *N,N*-dimethylthreonine.

Finally, we performed MS3 experiments to investigate whether the fragmentation patterns of the ion at *m/z* 130.086 ([M+H^+^]^+^) generated from a WT *P. aeruginosa* b-type glycan-modified peptide (**Fig. 2B**) and the methylated Type A-modified peptide from *C. difficile* (**Fig. 4A)** are the same. As shown in **Fig. 4B**, these spectra are identical, providing strong evidence that the hitherto unknown moiety in the *P. aeruginosa* b-type glycan is an *N,N*-dimethylthreonine.

## Discussion

The results from our study strengthen the notion that the b-type glycan in *P. aeruginosa* PAO1 and the Type A glycan in *C. difficile* strain 630 are very similar. Besides the difference in the core monosaccharide, the only difference in composition is the degree of *N*-methylation of the threonine as part of the glycan structure, i.e., in the b-type glycan in *P. aeruginosa* PAO1 it is dimethylated, while in the Type A glycan in *C. difficile* it is monomethylated.

The *m/z* of the *P. aeruginosa* b-type glycan-specific ion at *m/z* 356.110 that we observed in the MS/MS data is in agreement with previous data (15). Also in the earlier study, a fragment ion at *m/z* 130.1 on a low mass resolution instrument was observed in the MS/MS spectra of type-b glycan-modified peptides, but the identity was not elucidated (15). For our assignment, the data on the fragmentation of this ion, and comparison with methylated Type A from *C. difficile* was pivotal. Interestingly, several major fragments that we observed in the MS3 spectra, e.g. at *m/z* 70, 74, 84, 85, 86, 102 and 112, were observed in an MS/MS spectrum of *N,N*-dimethylthreonine analysed after CID fragmentation on a Q-TOF instrument (24). In the Type A glycan in *C. difficile,* the *N*-methylthreonine-phospho moiety is linked to the O-3 position on the GlcNAc. NMR analysis is necessary to determine whether the *N*,*N*-methylthreonine-phospho moiety in the *P. aeruginosa* b-type glycan is also in the same position on the rhamnose (14).

Our results confirm the b-type glycan site assignment on flagellin. Proteinase K digestion resulted in a set of b-type glycan-modified peptides with two b-type glycans and one of these (ASGIAS) had only two possible *O*-glycosyation sites, corresponding to Ser-191 and Ser-195 in the mature protein. Previously, these sites were determined based on the mass spectrometry analysis of a b-type glycan-modified peptide after beta-elimination (15).

Recently, we presented a revised model for the Type A biosynthesis in *C. difficile* strain 630 (19) based on a mass spectrometry-based proteomics experiment with mutant strains and bioinformatic analyses. The different steps in this model are schematically presented in **Fig. 5**.

**Figure 5:**
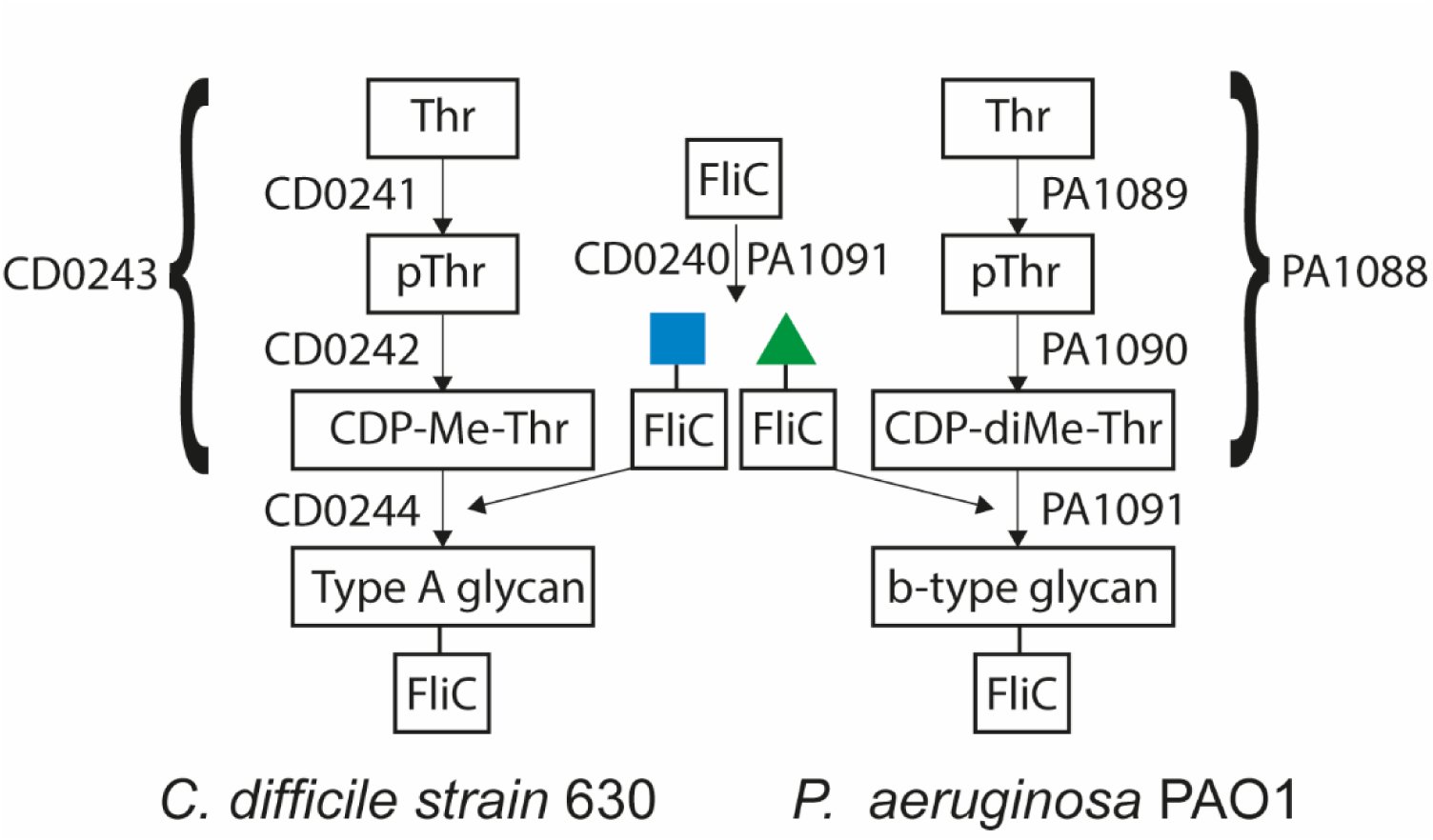
Schematic representation of the model for the biosynthesis of the flagellar glycans in *C. difficile* strain 630 and *P. aeruginosa* PAO1. For details about the individual steps and bioinformatic predictions of enzyme activities, see Claushuis et al. (19). The accolade indicates that it is unclear at what step the methylation occurs. Thr: threonine, pThr: phospho-threonine. Me: N-methyl, diMe: N,N-dimethyl

In this model, we predicted a novel biosynthetic intermediate, CDP-threonine (19), whose synthesis and subsequent role as a donor substrate is supposed to be controlled by CD0242 and CD0244, respectively. In *C. difficile*, the loss of CD0243 activity (a putative methyltransferase) partially resulted in a structure lacking the methyl group but our experimental approach did not allow us to determine the timing of the methylation event. The truncated b-type glycan in the putative methyltransferase (*pa1088*) *P. aeruginosa* mutant strain only consisted of the core rhamnose, in line with previous observations (15). Structures lacking methyl groups were not observed in this mutant strain, suggesting that methylation is a crucial, and early event in the b-type glycan biosynthetic pathway. Based on this finding, we offer the testable hypothesis that methylated intermediates are the preferred substrates for one or more of the enzymes in the biosynthesis routes. Hence, we here refine our model and postulate that CDP-*N*-methylthreonine and CDP-*N,N*-dimethylthreonine are the *in vivo* donor substrates of the reaction catalyzed by CD0244 and PA1091, respectively (**Fig. 5**). Of note, in addition to the structure lacking the methyl group, structures comprising only the core GlcNAc were also enriched in the *cd0243* mutant strain in *C. difficile* (19). We argued that this was due to polar effects in this strain but, assuming our model is correct, it might well be that the reaction with CDP-threonine as a donor substrate is suboptimal.

Because PA1091 encodes for both glycosyltransferase as well as phosphotransferase activity, it is unfortunately difficult to dissect the role of both activities independently. PA1090, a homolog of CD0242, is predicted to be responsible for the biosynthesis of CDP-(*N,N*-dimethyl)threonine. In line with this, we observed that the truncated b-type glycan in the corresponding mutant only contained the core monosaccharide. As expected, we also observed this phenotype in the *pa1089* mutant. Interestingly, previous data showed a mixture of WT structures and structures containing only the rhamnoses in a *pa1089* mutant (15). We have no explanation for this apparent discrepancy with our data other than that the mutant was generated using a different method.

In our quantitative proteomics data, we only observed a limited number of peptides corresponding to the enzymes involved in the b-type glycan biosynthesis, hampering reliable quantification. In *C. difficile* these proteins were readily identified and quantified (19), indicating that their overall cellular levels are higher. This might be related to the fact that *C. difficile* has multiple flagella per cell and more sites in FliC that are modified with the Type A glycan, whereas *P. aeruginosa* has only one flagellum per cell and only two modifications per FliC molecule. Ratio compression, known from TMT-based quantification methods, could also have played a role in our quantitative analyses. This is probably why the absence of the type-B modified peptide in the mutant strains was more apparent following IMAC purification. The minor residual signals that were observed in the *pa1088* and *pa1089* mutant strains in the IMAC data can largely be explained by impurities in the TMTpro labels; i.e., each label contains a small percentage of different isotopologues (TMT Reporter Ion Isotope Distributions for TMTpro 16plex batch WK334339, Thermo Fisher Scientific). The fact that we could use IMAC for affinity purification of b-type glycan-modified peptides demonstrated that the recently presented method (20) is more broadly applicable.

Both the innate and the adaptive immune response to flagellin is well-documented (25, 26). Interestingly, a *P. aeruginosa* flagella vaccine showed promising results in cystic fibrosis patients, highlighting the flagellin protein as a potential vaccine candidate (27). Our data may contribute to future vaccine designs, even though recent data suggested that the b-type glycan is less important than the a-type glycan for the induction of protective antibodies (28). Moreover, further understanding and characterization of the enzymes involved in the b-type glycan biosynthesis may lead to the development of potential inhibitors, some of which may even be able to target *P. aeruginosa* as well as *C. difficile* strains.

## Experimental procedures

### Bacterial strains and culturing conditions

The wildtype *P. aeruginosa* PAO1 strain was a generous gift from prof. A. Briegel from the Institute of Biology Leiden (IBL) in Leiden, the Netherlands. The PAO1 mutant strains were ordered from the Salipante Lab at the University of Washington, USA (21, 22). Information about the mutant strains can be found in Supporting Table S1. Cells were plated on Luria-Bertani (LB) agar plates. Three colonies per plate (representing three biological replicates per strain) were inoculated in 5 mL LB medium and grown for 16 h at 37°C, while rotating at 180 rpm. *C. difficile* strain 630 Δ*erm* was cultured as described previously (19).

### Protein extraction, reduction alkylation, digestion and TMT-labeling

Following cell culturing, cells were pelleted by centrifugation (3220xg, 10 min, 4 °C), resuspended in 5mL of ice-cold PBS, and centrifuged again. The washing step was repeated once more. After the last wash, pellets were resuspended in 1 mL of ST lysis buffer (5% SDS, 0.1 M Tris-HCl pH 7.5) and incubated on ice for 20 minutes. Then, cells were lysed using sonication for 30 seconds, followed by 30 seconds of cooling on ice. This was repeated five times. After sonication, samples were centrifuged at 10000 x g for 15 minutes at room temperature (RT), and the supernatant was collected. Protein levels were quantified using a BCA assay (Pierce Rapid Gold BCA Protein Assay). For reduction and alkylation of cysteines, 1 µL 0.5 M tris(2-carboxyethyl)phosphine, 3 µL 0.3 M iodoacetamide, and 2 µL 0.5 M dithiothreitol were sequentially added to a protein solution (100 µg in 100 µL ST buffer), with each step incubated for 30 minutes at RT. Then proteins were precipitated by sequentially adding 400 µL methanol, 100 µL chloroform, and 300 µL water, with vortexing after each step. Following centrifugation at max speed in an Eppendorf 5424 R centrifuge, the protein pellet at the interface was collected and washed three times with 500 µL methanol. The pellet was then air-dried at 37 °C.

For Proteinase K (Merck) digestion, a protein pellet corresponding to 100 µg protein was reconstituted in 100 µL 100 mM ammonia solution containing 4 µg Proteinase K, after which samples were incubated overnight at 37°C. For the combined trypsin (Worthington Biochemical)/chymotrypsin (Worthington Biochemical) digestion, a protein pellet corresponding to 100 µg protein was resuspended in 100 μl 40 mM HEPES pH 8.0 containing 2 μg trypsin and 2 μg chymotrypsin, and incubated overnight at 37°C, after which an additional 2 μg of each enzyme was added. Samples were incubated for two hours. Tandem Mass Tagging (TMT, Thermo) and subsequent peptide fractionation (12 fractions) were performed as described previously (19) using 10 μg of tryptic+chymotryptic peptides from each strain. For the overview of the labels for each strain, see **Table S2**.

### Fe^3+^-Immobilized Metal Affinity Purification (IMAC)

Before IMAC, 200 µg of peptides were desalted using solid phase extraction (SPE) on an HLB Oasis 1 cc cartridge (Waters). First, the cartridge was activated using 1 ml 10/90 (v/v) H2O/acetonitrile (AcN) and then equilibrated using three times 1 ml 0.1% trifluoroacetic acid (TFA). After sample loading, the cartridge was washed three times with 1 ml 0.1% TFA. Peptides were eluted in 400 μl 30/70/0.1 AcN/H_2_O/TFA and freeze-dried. A Fe(III)-NTA cartridge (Agilent) was prepared by priming it with 250 μl 0.1% TFA in AcN. The cartridge was washed three times with 250 μl 0.1% TFA in AcN. The freeze-dried eluate from the SPE was dissolved in 200 µL 30/70/0.1 (v/v/v) AcN/H_2_O/TFA and loaded on the IMAC cartridge, and the cartridge was washed three times with 250 μl 0.1% TFA in AcN. Peptides were eluted with 25 μl 1% ammonia directly into 25 μl 10% formic acid (FA), and freeze dried.

### Dimethylation of *C. difficile* tryptic peptides

A tryptic digest of *C. difficile* strain 630Δerm proteins was generated as described previously (19). Subsequently, 50 μg of tryptic peptides were diluted in 0.1% FA to a final volume of 1 ml and applied to a HLB Oasis 1 cc cartridge (Waters) which was activated using 1 ml 10/90 (v/v) H2O/ACN and equilibrated using three times 1 ml 0.1% formic acid (FA). After washing three times with 1 mL 0.1% FA, the dimethylation labelling mixture (4.5 mg sodium phosphate cyanoborohydride (NaCNBH3), 14 μl formaldehyde (CH2O) 37%/2.5 ml sodium phosphate buffer pH 7.5) was added for 5 minutes, applying 0.5 ml at 1 minute-intervals of incubation. Then, the cartridge was washed three times with 0.1% FA. Finally, labelled peptides were eluted with 400 μl of an 80/20/0.1 (v/v/v) AcN/water/FA solution and freeze-dried.

### Analysis of TMT-labeled *P. aeruginosa* flagellin peptides

For the data dependent analyses, the PAO1 TMT-labeled peptides were dissolved in 0.1% FA and analyzed by online C18 nanoHPLC MS/MS using an Ultimate3000nano gradient HPLC system (Thermo, Bremen, Germany) coupled with an Exploris480 mass spectrometer (Thermo). Fractions were injected onto a cartridge precolumn (300 μm × 5 mm, C18 PepMap, 5 μm, 100 A, (100/0.1 water/(FA) v/v) at a flow of 10 μl/min for 3 minutes (Thermo, Bremen, Germany) and eluted via a homemade analytical nano-HPLC column (30 cm × 75 μm; Reprosil-Pur C18-AQ 1.9 μm, 120 A (Dr. Maisch, Ammerbuch, Germany) at a flow of 250 nL/min. The gradient was run from 2% to 36% solvent B (20/80/0.1 water/acetonitrile/formic acid (FA) v/v/v) in 120 min. The temperature of the nano-HPLC column was set to 50°C (Sonation GmbH, Biberach, Germany). The nano-HPLC column was drawn to a tip of ∼10 μm and acted as the electrospray needle of the MS source. The mass spectrometer was operated in data-dependent MS/MS mode for a cycle time of 3 seconds, with a normalized HCD collision energy of 36 % and recording of the MS2 spectrum in the Orbitrap. In the master scan (MS1) the resolution was 120,000, the scan range *m/z* 350-1600, at a Standard AGC target with a maximum fill time of 50 ms. Dynamic exclusion after n=1 with an exclusion duration of 45 s and a mass tolerance of 10 ppm. Charge states 2-5 were included. For MS2 precursors were isolated with the quadrupole with an isolation width of 1.2 Da. The first mass was set to 110 Da and the MS2 scan resolution was 30,000. All raw data were converted to peak lists using Thermo Proteome Discoverer 2.4.1.15. and searched against the *P. aeruginosa* PAO1 database (strain ATCC 15692, downloaded from Uniprot on April 13, 2020, number of entries: 5564) using Mascot v. 2.2.07. Trypsin+chymotrypsin (C-term FKLRWY, not before P) were selected as enzyme specificity with a maximum of two missed cleavages. Mass tolerances of 10 ppm and 0.02 Da for precursor and fragment ions were used, respectively. TMTPro (K), TMTPro (N-term), and Carbamidomethyl (C) were selected as fixed modifications. Methionine oxidation, acetylation (Protein N-term) and wildtype b-type glycan (neutral mass 355.103 Da) were selected as variable modifications. Peptides with an FDR < 1% based on Percolator (23) were accepted. Quantification of the data was performed using the reporter ion intensities of TMTpro labels to compare relative peptide abundance across samples.

The IMAC purified TMT-labeled peptides were analyzed by online C18 nanoHPLC MS/MS using an Easy nLC 1200 gradient HPLC system (Thermo, Bremen, Germany), and an Orbitrap Fusion LUMOS mass spectrometer (Thermo). The LC gradient was similar as described above. Instead of a data dependent analysis, MS/MS was performed on the predefined precursor ion at *m/z* 1116.318 ([M+4H^+^]^4+^), corresponding to the tryptic + chymotryptic *Pseudomonas aeruginosa* TMT-labeled FliC peptide QVGSNGAGTVASVAGTATASGIASGTVNLVGGGQVK with two Type B modifications. The first mass was set to 110 Da and the MS2 scan resolution was 30,000. Quantification of this peptide was again performed using the reporter ion intensities of TMTpro labels.

### Mass spectrometry of *P. aeruginosa* ProtK flagellin peptides and methylated Type A-modified tryptic flagellin peptides from *C. difficile*

For the MS analysis of IMAC purified *P. aeruginosa* ProtK peptides, samples were dissolved in 0.1% FA and analyzed by online C18 nanoHPLC MS/MS using an an Easy nLC 1200 gradient HPLC system (Thermo, Bremen, Germany), and an Orbitrap Fusion LUMOS mass spectrometer (Thermo). Fractions were injected onto a homemade precolumn (100 μm × 15 mm; Reprosil-Pur C18-AQ 3 μm, Dr Maisch, Ammerbuch, Germany) and eluted via a homemade analytical nano-HPLC column (30 cm × 75 μm; Reprosil-Pur C18-AQ 1.9 μm). The analytical column temperature was maintained at 50 °C with a PRSO-V2 column oven (Sonation, Biberach, Germany). The gradient was run from 2% to 40% solvent B (20/80/0.1 water/acetonitrile/formic acid (FA) v/v) in 120 min. The nano-HPLC column was drawn to a tip of ∼5 μm and acted as the electrospray needle of the MS source. The MS spectrum was recorded in the Orbitrap (resolution 120,000; *m/z* range 400–1500; maximum injection time 50 ms). Dynamic exclusion was after n = 1 with an exclusion duration of 15 s with a mass tolerance of 10 ppm. Charge states 1–5 were included.

For MS/MS(/MS) analysis, samples were re-analyzed on the above system using a shorter gradient (2-40%B in 60 min). Fragmentation was performed on the predefined precursor ion at *m/z* 737.797 ([M+2H^+^]^2+^), corresponding to the flagellin peptide TASGIASGT with two b-type glycans. For the subsequent MS3 experiments, fragment ions at *m/z* 356.110 and 130.086. The same MS/MS(/MS) method was used for the methylated Type A-modified tryptic peptides from *C. difficile* flagellin with predefined precursor and fragment ions at *m/*z 751.871 and 130.086, respectively.

## Supporting information

This article contains supporting information

## Acknowledgements

We thank prof. Ariane Briegel (Leiden University) for providing the WT PAO1 strain.

## Author contributions

P.J.H. conceptualization; P.J.H., L.v.H. formal analysis; L.v.H., J.v.A., A.H.de.R. B.C. investigation; P.J.H., B.C. methodology; P.A.v.V. validation; P.J.H., L.v.H., visualization; P.J.H., writing–original draft; P.J.H., J.C., P.A.v.V., B.C., W.K.S. writing–review & editing; P.A.v.V., W.K.S., J.C. resources; P.J.H., P.A.v.V., J.C., P.J.H. supervision; P.J.H. data curation; P.A.v.V. funding acquisition; P.J.H., J.C., P.A.v.V. project administration;

