## Supplemental figures and tables for "Elucidation of the glycan structure of the b-type flagellin of *Pseudomonas aeruginosa* PAO1"

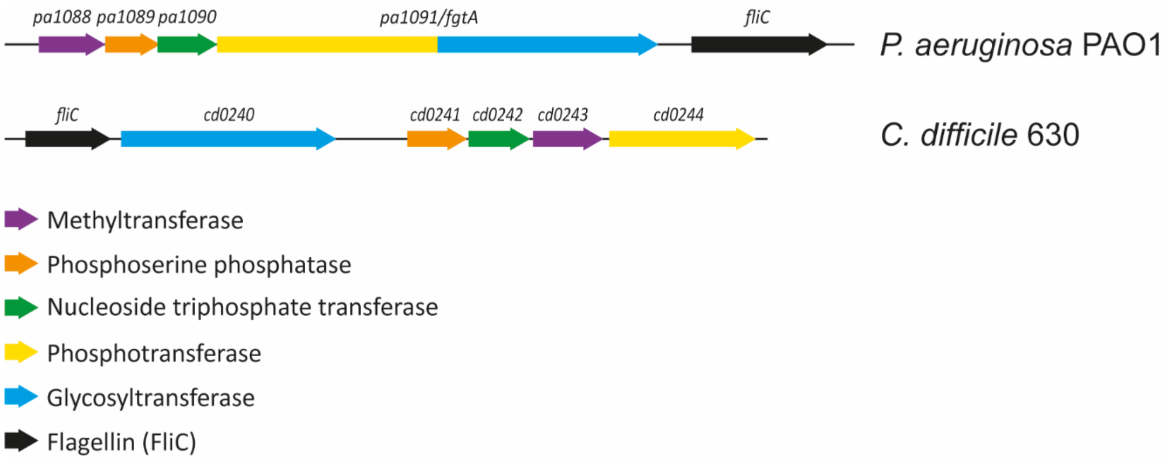


**Fig. S1**: **Flagellin glycosylation biosynthetic gene clusters in *Pseudomonas aeruginosa* PAO1 and *Clostridioides difficile* strain 630*Δerm*.**

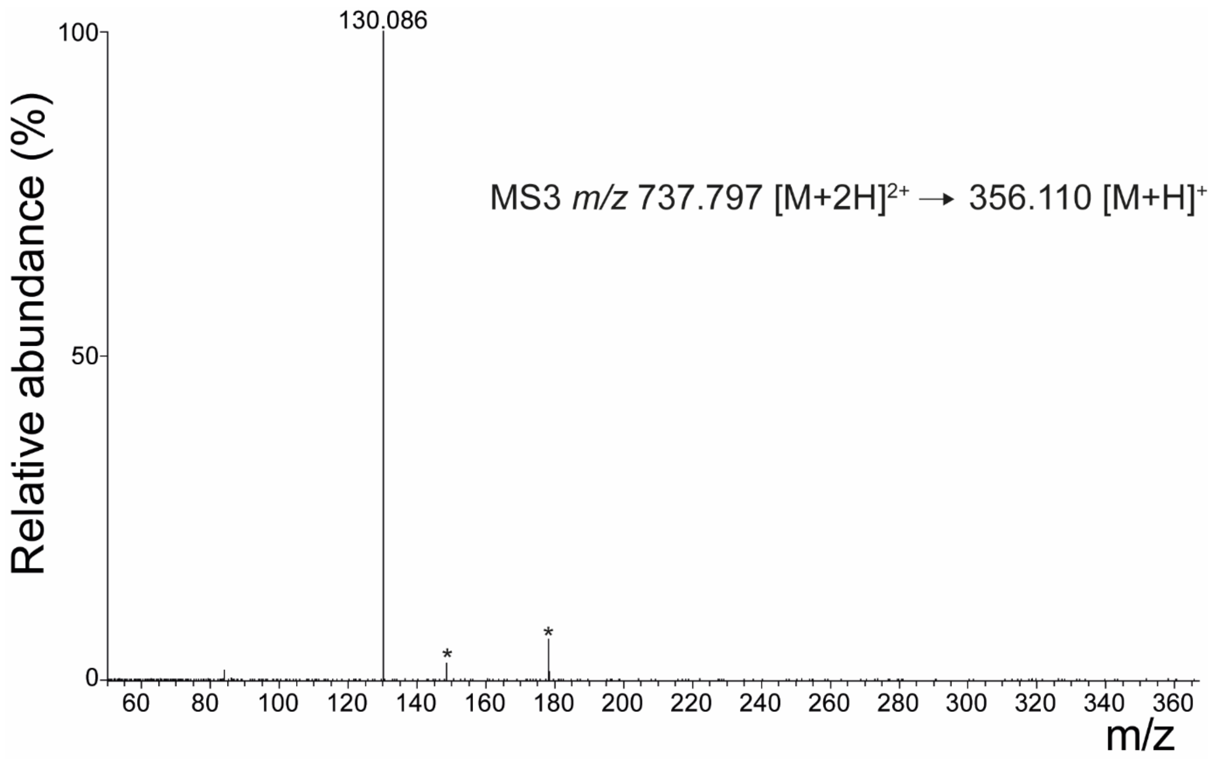


**Fig. S2**: **MS3 fragmentation of the Type A-specific fragment ion at *m/z* 356.110 as observed in Fig. 3A**. The signals indicated with an asterisk (*) are background signals of unknown origin that were observed in all fragmentation spectra.

```
>tr|Q9I4N8|Q9I4N8_PSEAE Flagellar glycosyl transferase, FgtA OS=Pseudomonas
aeruginosa (strain ATCC 15692 / DSM 22644 / CIP 104116 / JCM 14847 / LMG
12228 / 1C / PRS 101 / PAO1) OX=208964 GN=fgtA PE=4 SV=1
MIEDSVSEPDRRDGGDAGKRLAWLQRQVLPVLEEGGAVWVSGLDGAPFAGGAHKVVEASV
AGELPGNERFRLACLGGAVGGVADDWQAMRLLLRAVESLEDRGWLLLEEALSVSAAGACR
SPQAQARLALSLGLRQVAELRLGAPDDGGRQRVLQLFRQDLAVARMQRYSGLRVACYGNM
PFHYRSLRPLAECFEDSLLSLDIDEVMAWKPDVIAVADGWSVEFWRDYCDAHNVLLVGMR
HGSVTRYGFAEGTYRYADYLCGSAWDIDDTLASSVMPRNGFLLTGNAWCDEVFRLPARTP
AENAPTILFAPTYNPEISAAVHLGERVVALIRKVYPASRIIIKPHPAIVQHEHAFVSDKD
LFRDLMKLWREQSRTDPLVTLVDDPEASIAASFAEADILLADRSSLIFEFMTLDRPILLF
SREQRIARWAYNPEAPGNAWRDIGLEFADDEQLLDLLANAFTRHAESRDTQENRTQQLYG
RFRDGRSVQRVAAAIAEAPRLQIVLDARAAADGGQRLAEHYAACFAHRRIDLIAPSGAAL
PADVRRFDSWRAWCDAEAATLERRASAVLLVEDDGQLLPGSAHQVSALLPAIAMGQLRHA
RLALEEREAGPAGETDWSARRLHRLLARLQPKPAWRLLAPALLREELADLPDEAEQPLWL
RAVAAAEAPVEWPVGQLNLFAEEGLLRQGDVHMLCARARLRAVPCVSGLPLRQRQVVLQV
TPFASQLYDRFPFQLRVCLNGQLFSRVSVEDARAKSLVLPFRGDERGGMTIELECDGSYP
AFEGVSTPVSLLLQTAFQPAPGDYCAALAAGAASPEAGEDLRSEYETWLHGREVEPARYR
PFIDALRRDSRIEVLVLAEEAEDADLQRSLASIDGQALPAWRTRILGRAPAFARKGLAWL
AEGGSAAERINLAAAASDADWLIVIHAGDELARSALLLLAEKIRTETALLCCYSDEDHVC
DGRYEAPLLKPDFNLDLLRSYPYCGRSLAFQRAALLAQGGLQEGFGDLALQDFMFRLAER
EGLDRIGHLAEVLYHSARAFGEWLASAAVRPFIASVVDEHLNRLGVPHRIEPGRLAVINR
IAYDYPGTPAVSLLLPVGDSLSALQRSVESFLENTDYPSYELLLVASGPLAPDMAAWLEA
VQGLGSEQIRVLSPQASSLAGCLNLCVAEARGEFLLSLGAGVVALRPDWLRELLNHGRRP
EVGAVGGKLLGLDGTIREAGLVLGLGGTAGRAFAGEAGDSAGYMHRLLVVQNHTALSASC
LLFKRSLHDELGGFDENDFAHGHADVDFSLRARQLGYLSVWTPYAILAQNGNVELPSVEA
DESLYRRWLPALARDPAYNRNLSLEGAGFALERPEVPAWQPLFGRSPLPRVLAHPADPYG
CGHYRVRQPFRALHDAGLLDGMLSESLLQPVALERLEVDSVILQRQISEEQLRAISRMRS
FNRAFRVYELDDYLPELPLKSLHRAEMPGDIRQILGRALGLADRFVVSTEPLAEAFRRMH
GDIRVVPNRLPLPWWRDLSSRRRDAERPRVGWAGGIGHGGDLEVIAEVVRELADEVDWVF
FGFCPDALRPYVREYHPGVEIERYPAYLASLDLDLALAPLEQNRFNECKSNLRLLEYGVL
GFPVICSDVLCYRDSLPVTRVKNRSRDWLEAIRAHLADADANAAAGAQLREAVRRDWMLE
GAHLEAWAAAWLPD
```

**Fig. S3**: **Relative quantification of FgtA (PA1091) peptides in the pa1091 mutant strain based on a quantitative proteomics experiment using TMT.** Peptides colored in red had a ratio of pa1091 mutant/WT below 0.7 (0.27-0.65, median 0.50, n=8). Peptides colored in yellow had a ratio of pa1091 mutant/WT above 0.7 (0.78-1.39, median 0.92, n=14).

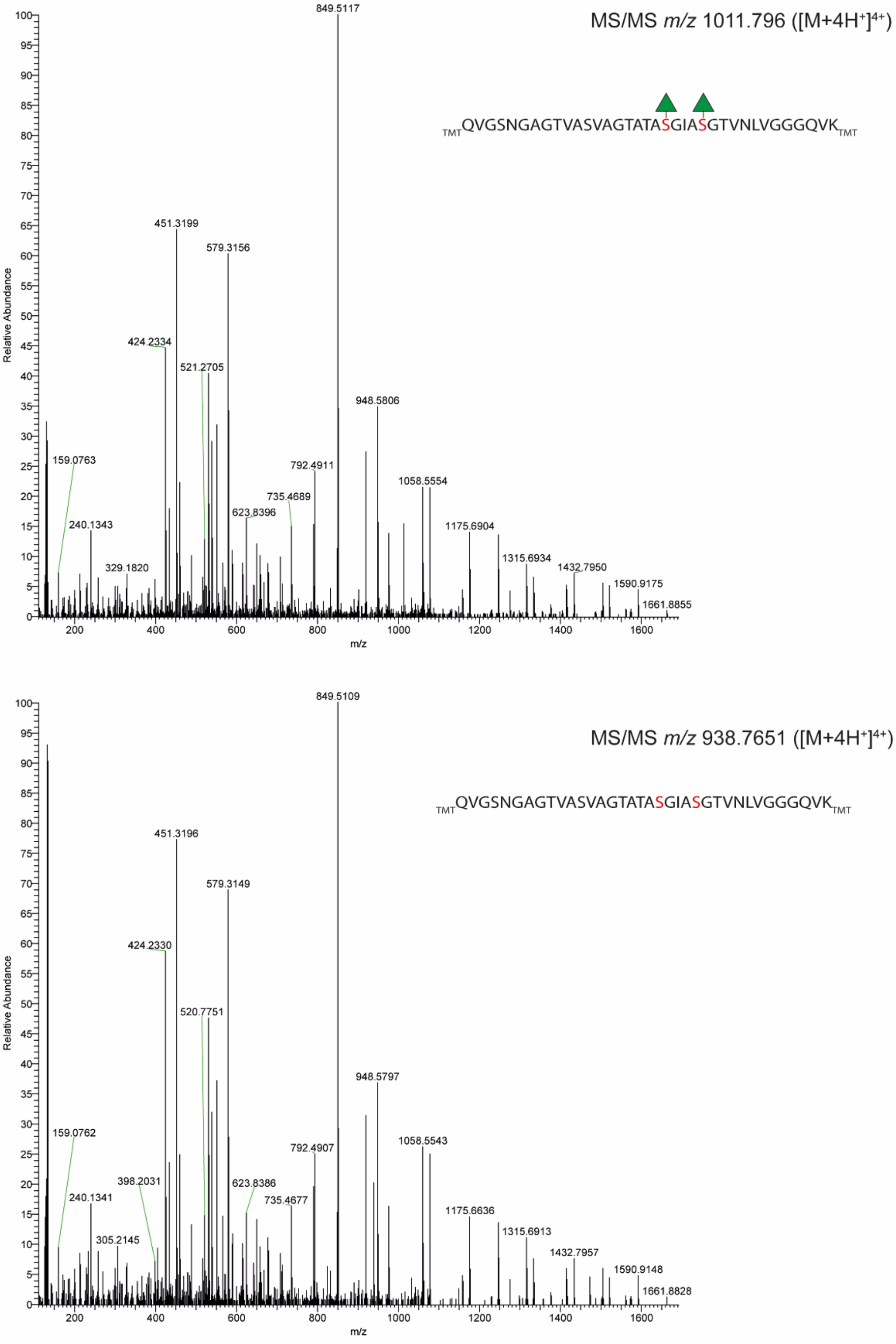


**Fig. S4**: **MS/MS spectra of the TMT-labeled flagellin tryptic+chymotryptic peptide QVGSNGAGTVASVAGTATASGIASGTVNLVGGGQVK with two rhamnoses (upper panel) or non-glycosylated (lower panel)**.

**Table S1**: Overview of the *P. aeruginosa* PAO1 strains used in this study. *For more information about the transposon mutants see Jacobs et al. 2003, PNAS 100:14339 and Held et al. 2012, J. Bacteriol. 194:6387. *: strain ID from the Salipante lab (https://sites.google.com/uw.edu/salipante-lab).*

| Strain | Original strain ID* | Genotype | Locus tag | Description |
|---|---|---|---|---|
| BC107 | n.a. | WT PAO1 | n.a. | WT |
| BC108 | PW2962 | PA1088-C10::ISLacZ/hah | PA1088 | *pa1088* mutant |
| BC111 | PW2965 | PA1089-F11::ISphoA/hah | PA1089 | *pa1089* mutant |
| BC112 | PW2967 | PA1090-A07::ISphoA/hah | PA1090 | *pa1090* mutant |
| BC115 | PW2969 | PA1091-A02::ISphoA/hah | PA1091 | *pa1091* mutant |

**Table S2.** *Overview of the TMTpro labels per strain.*

| **TMTpro label** | **Strain** | **Genotype** | **Description** |
|---|---|---|---|
| 126 | BC107 | WT | WT |
| 127N | BC107 | WT | WT |
| 127C | BC107 | WT | WT |
| 128N | BC108 | PA1088-E05::ISLacZ/hah | *pa1088* mutant |
| 128C | BC108 | PA1088-E05::ISLacZ/hah | *pa1088* mutant |
| 129N | BC108 | PA1088-E05::ISLacZ/hah | *pa1088* mutant |
| 129C | BC111 | PA1089-B10::ISphoA/hah | *pa1089* mutant |
| 130N | BC111 | PA1089-B10::ISphoA/hah | *pa1089* mutant |
| 130C | BC111 | PA1089-B10::ISphoA/hah | *pa1089* mutant |
| 131N | BC112 | PA1090-B10::ISphoA/hah | *pa1090* mutant |
| 131C | BC112 | PA1090-B10::ISphoA/hah | *pa1090* mutant |
| 132N | BC112 | PA1090-B10::ISphoA/hah | *pa1090* mutant |
| 132C | BC115 | PA1091-B10::ISphoA/hah | *pa1091* mutant |
| 133N | BC115 | PA1091-B10::ISphoA/hah | *pa1091* mutant |
| 133C | BC115 | PA1091-B10::ISphoA/hah | *pa1091* mutant |

**Table S3:** Relative quantification of PA1088-PA1089 in the corresponding mutant strains (BC108, BC111, BC112 and BC115)

| Gene | Protein | Uniprot ID | Coverage (%) | #peptides | #PSMs | #quant peptides | Mutant/WT |
|---|---|---|---|---|---|---|---|
| *pa1088* | PA1088 | Q9I4P1 | 13 | 4 | 4 | 4 | 0.5 (BC108/BC107) |
| *pa1089* | PA1089 | Q9I4P0 | 27 | 4 | 4 | 2 | 0.8 (BC111/BC107) |
| *pa1090* | PA1090 | Q9I4N9 | 16 | 2 | 2 | 1 | 0.5 (BC112/BC107) |
| *pa1091* | FgtA (PA1091) | Q9I4N8 | 23 | 36 | 42 | 21 | 0.7 (BC115/BC107) |
